# The architecture of Cidec-mediated interfaces between lipid droplets

**DOI:** 10.1101/2021.03.23.436555

**Authors:** Iva Ganeva, Koini Lim, Jerome Boulanger, Patrick C. Hoffmann, David B. Savage, Wanda Kukulski

## Abstract

Lipid droplets (LDs) are intracellular organelles responsible for storing surplus energy as neutral lipids. Their size and number vary enormously (1). In white adipocytes, they reach up to 100 *µ*m in size, occupying >90% of the cell. Cidec, which is strictly required for the formation of such large LDs, is concentrated at interfaces between adjacent LDs and facilitates the directional flux of neutral lipids from the smaller to the larger LD (2-7). However, the mechanism of lipid transfer is unclear, in part because the architecture of interfaces between LDs has remained elusive. Here we visualised interfaces between LDs by electron cryo-tomography (cryo-ET) and analysed the kinetics of lipid transfer by quantitative live fluorescence microscopy (FM). We show that transfer occurs through closely apposed intact monolayers, is slowed down by increasing the distance between the monolayers and follows exponential kinetics suggesting a pressure-driven mechanism. We thus propose that unique architectural features of LD-LD interfaces are mechanistic determinants of neutral lipid transfer.

## Main Text

White adipocytes, which are responsible for regulated fat storage and release in vertebrates, feature a single large LD filling up almost their entire cytoplasm. This distinctive morphology is a key factor in optimising energy storage capacity and reducing the relative surface area, thereby enabling tight regulation of lipid storage and release from the droplet (1). Excessive or deficient lipid storage are hallmarks of several serious and highly prevalent metabolic diseases such as diabetes, non-alcoholic fatty liver disease and atherosclerosis. Both human and mouse genetic studies suggest that Cidec is strictly required for the formation of unilocular white adipocytes (4-6).

To emulate the requirement for Cidec, we first used *Cidec* null mouse embryonic fibroblast (MEF)-derived adipocytes (4) and expressed Cidec-EGFP during adipogenic differentiation using a lentiviral Tet-ON inducible system. Upon expression, Cidec-EGFP was targeted to the surface of LDs and enriched at interfaces between LDs (Fig. 1a). The volume of LDs increased as compared to LDs in cells that were not induced (Fig. 1a)(n=2 repeats, mean -Dox 4.8 *µ*m^3^, mean +Dox: 59.1 *µ*m^3^). Expression of cytosolic EGFP had no effect on LDs (Suppl. Fig. S1a). These results show that induced expression of Cidec-EGFP in MEF-derived *Cidec* null adipocytes recapitulates the localisation and activity of Cidec on LDs, consistent with previous observations in other cell types (3, 8-10).

**Figure 1:**
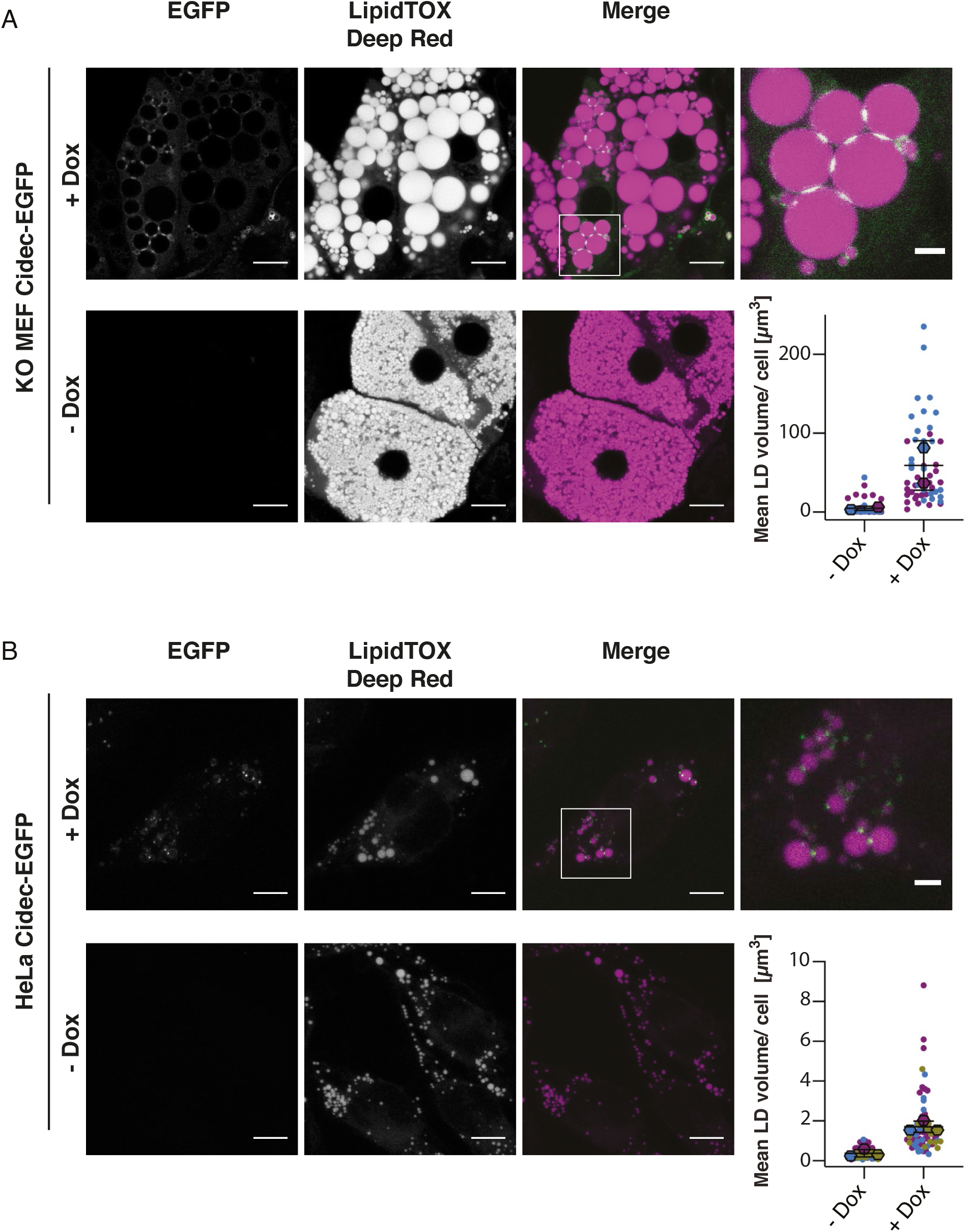
Induced expression of Cidec-EGFP increases LD sizes in Cidec null MEF-derived adipocytes and in HeLa cells. **A)** Fluorescence microscopy of fixed Cidec null MEF-derived adipocytes inducibly expressing Cidec-EGFP (green). LDs were labelled with LipidTOX Deep Red dye (magenta). Cidec null MEFs were differentiated into mature adipocytes over the course of 10 days in the continuous presence (upper panel) or absence (lower panel) of Doxycycline. Scale bars: 10 *µ*m. The last image in the upper panel shows enlargement of the boxed region. Scale bar: 2.5 *µ*m. The plot shows mean LD volumes per cell. Means of individual experimental repeats are represented by diamond symbols, individual cells are represented by dots, colour coded according to experiment. Black lines correspond to the mean of two experiments with standard deviation (SD) (n=2 repeats each analysing 25 cells per conditions, -Dox: 4.75 *µ*m^3^, SD 2.46 *µ*m^3^; +Dox: 59.08 *µ*m^3^, SD 31.49 *µ*m^3^). **B)** Fluorescence microscopy of fixed HeLa cells inducibly expressing Cidec-EGFP (green). LDs were stained with LipidTOX Deep Red dye (magenta). The upper panel represents HeLa cells induced with Doxycycline for Cidec-EGFP expression. The lower panel shows HeLa cells in the absence of Doxycycline induction. Cells were fixed 24 hours after the addition of oleic acid and Doxycycline. Scale bars: 10 *µ*m. The last image in the upper panel shows enlargement of the boxed region. Scale bar: 2.5 *µ*m. The plot shows mean LD volumes per cell. Means of individual experimental repeats are represented by diamond symbols, individual cells are represented by dots, colour coded according to experiment. Black lines correspond to the mean of three experiments with standard deviation (SD) (n=3 repeats each analysing 15-32 cells per condition, -Dox: 0.38 *µ*m^3^, SD 0.18 *µ*m^3^; +Dox: 1.71 *µ*m^3^, SD 0.28 *µ*m^3^).

We next wished to exploit this experimental paradigm to investigate the architecture of the interface between LDs by cellular cryo-ET. Although the close interaction between LDs has been described as a LD contact site (3), some mechanistic models suggest that Cidec facilitates the formation of fusion pores between the two monolayers (8, 11). It is however unclear whether fusion of the two monolayers occurs, or whether the LD interface is instead reminiscent of organelle contact sites formed between topologically separate membranes (12). To address this question, we transferred the Tet-ON inducible expression of Cidec-EGFP to HeLa cells, for which cryo-ET workflows are well established (13, 14). Expression of Cidec in heterologous cells that do not express any adipocyte-specific proteins induces LD enlargement, indicating that Cidec alone is sufficient for this process (3, 9, 10). We ‘fed’ the HeLa cells with oleic acid for 24 hours to trigger LD formation. Although LDs in HeLa cells are considerably smaller than in MEF-derived adipocytes, Cidec-EGP expression increased LD volumes (Fig. 1b) (n=3 repeats, mean -Dox: 0.38 *µ*m^3^, mean +Dox: 1.71 *µ*m^3^, P=0.0024). Expression of cytosolic EGFP had no effect (Suppl. Fig. S1b). These results verify that expression of Cidec-EGFP in HeLa cells is sufficient to replicate its function in LD enlargement.

We thus vitrified HeLa cells on cryo-EM grids after feeding them with oleic acid and inducing Cidec-EGFP expression. To identify Cidec-EGFP enriched at LD interfaces, we imaged the vitrified cells by FM at cryogenic temperatures (cryo-FM) (Suppl. Fig. S2a and b). Subsequently, we thinned corresponding cell regions by cryogenic focused ion beam milling (cryo-FIB milling) (Suppl. Fig 2c) and subjected them to cryo-ET (13). This approach allowed us to visualize LD interfaces marked by Cidec-EGFP in a near-native state and at high resolution. In the resulting tomograms, we could readily identify LDs based on their dense, amorphous core consisting of neutral lipids including triacylglycerol (TAG) (Fig. 2) (15). The surrounding monolayers were resolved as single dark lines, as compared to the typical two lines visible for bilayers (Suppl. Fig. 2g). Where two LDs were closely apposed, we often observed deformations of the otherwise nearly spherical shape of the LDs. These large-scale morphologies varied among the interfaces we imaged, and corresponded to either minimal deformation (Fig. 2a, Movie 1), flattening of both LDs (Fig. 2b, Movie 2), one LD locally protruding and inducing an indentation in the other LD (Fig. 2c, Movie 3), or the smaller LD locally imposing its curvature by inducing an indentation in the larger LD (Fig. 2d, Movie 4).

**Figure 2:**
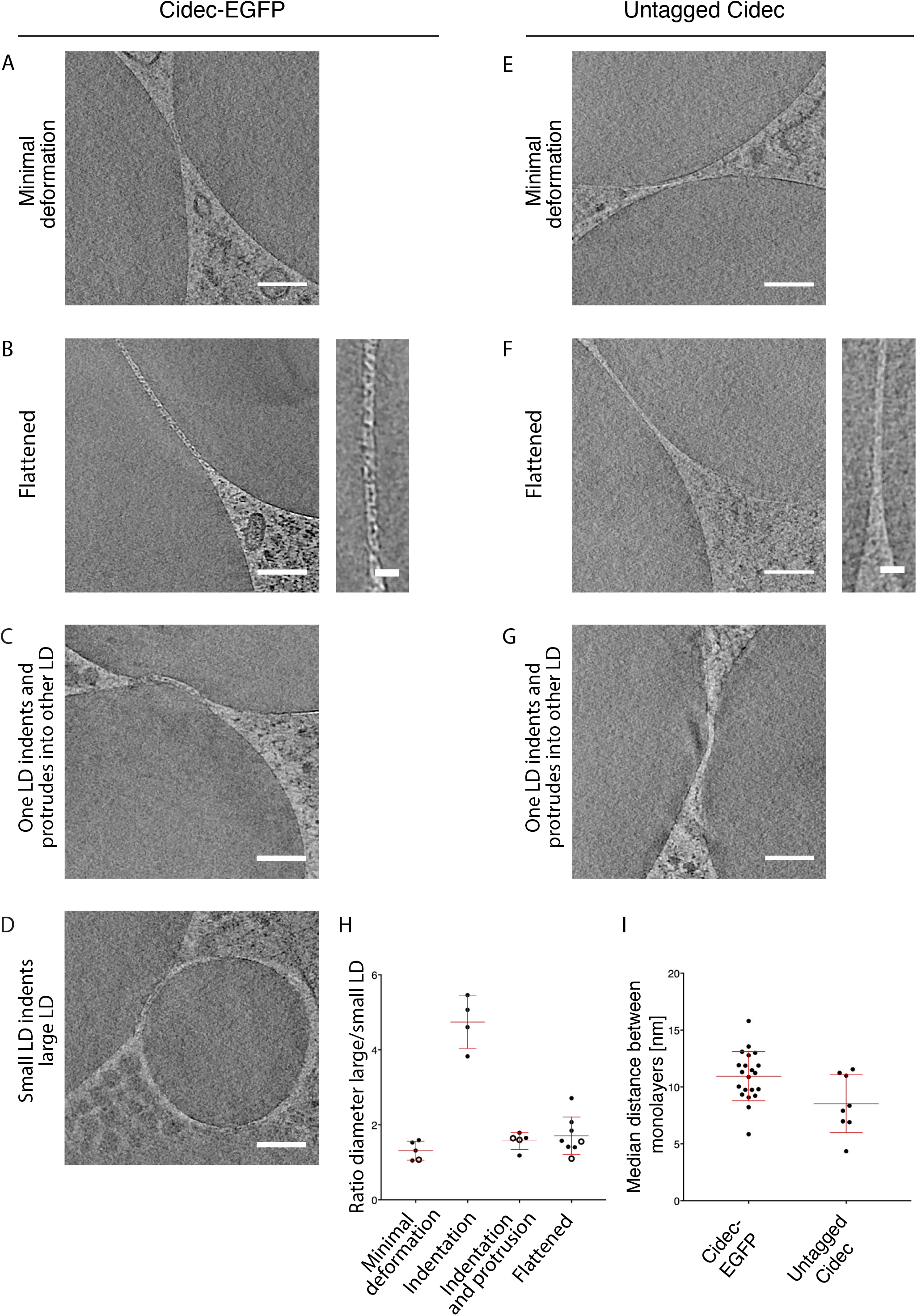
Electron cryo-tomography of LD interfaces in HeLa cells expressing Cidec-EGFP or Cidec. Virtual slices of tomograms acquired at areas where LDs are in close proximity. HeLa cells inducibly expressing Cidec-EGFP **(A-D)** or untagged Cidec **(E-G)** were vitrified, screened for LD interfaces by cryo-FM, thinned by cryo-FIB milling and imaged by cryo-ET. The shape of the LDs is deformed where two LDs are closely apposed. The different observed morphologies are classified as follows: **A) and E)** minimal deformation of the monolayers; **B) and F)** flattening of both LDs forming an interface; **C) and G)** protrusion of one LD into the other LD, resulting in an indentation. **D)** indentation of the larger LD caused by the smaller LD. Insets in B and F show a higher magnification view of the monolayers at the interface. **H)** The different interface morphologies plotted as a ratio of the diameter of the larger LD to the diameter of the smaller LD. Filled black dots indicate Cidec-EGFP data, and empty circles indicate data from untagged Cidec. Red bars represent mean with standard deviation (SD) (minimal deformation: 1.31, SD 0.25; indentation: 4.74, SD 0.70; indentation and protrusion: 1.57, SD 0.23 ; flattened: 1.71, SD 0.50). Only LD pairs where the sample thickness allowed a reliable diameter measurement (see Materials and Methods) were included in this quantification. **I)** Median distances between each LD pair, for Cidec-EGFP and untagged Cidec. Red bars represent the mean with SD (Cidec-EGFP: 10.94 nm, SD 2.16 nm; Cidec: 8.53 nm, SD 2.55 nm). Scale bars in A-G: 100 nm; scale bars in insets: 25 nm.

In all cases, the two phospholipid monolayers appeared as separate entities with no continuity between them. We determined that the mean distance between closely apposed monolayers forming LD-LD interfaces was 10.9 nm (n=21, SD 2.16 nm) (see Materials and Methods). Furthermore, we observed a dense layer of material between the monolayers, likely corresponding to proteins (Fig. 2b, enlargement). Given the dimensions of the interface and the dense protein packing, we hypothesised that the bulkiness of the Cidec-EGFP construct might influence the architecture of the interface.

To address the influence of the Cidec-EGFP construct size, we generated a HeLa cell line in which we inducibly expressed untagged Cidec. Using cryo-FM, we identified in these cells LipidTOX-labeled LDs closely apposed to each other, hence likely engaged in a LD-LD interface (Suppl. Fig. S2d and e). We targeted these areas by cryo FIB-milling and subsequent cryo-ET (Suppl. Fig. S2f). We found that the overall architecture of LD interfaces was consistent between HeLa cells expressing Cidec-EGFP and untagged Cidec. As for Cidec-EGFP, the two apposed monolayers were visible as two separate entities in cells expressing untagged Cidec (Fig. 2e-g, Movies 5-7). We did not observe continuity between the monolayers of the two LDs, indicating that at all interfaces we imaged, there was no fusion of monolayers. This finding implies that TAG transfer occurs through two intact phospholipid monolayers, rather than a fusion pore.

The LD pairs in cells expressing untagged Cidec displayed similar large-scale morphologies to those observed for Cidec-EGFP, indicating that these deformations are inherent to LD-LD interfaces (Fig. 2e-g, Movie 5-7). We thus analysed whether the different morphologies correlated with the ratio of diameters of the LDs engaged in the interface, or with the absolute sizes of the LDs. Minimal deformation, flattening, and protruding LDs, were morphologies found at ratios of large-to-small LD diameter between 1 and 2.7 (Fig. 2h). In contrast, small undeformed LDs forming an indentation in large LDs were found when the diameter ratio was between 3.8 and 5.5. This type of morphology was exclusively associated with LDs of less than 200 nm in diameter, and such small LDs were never associated with other morphologies. These data suggest that at diameters of less than 200 nm, LDs maintain a spherical shape when interacting with large LDs and impose their curvature locally on the interacting LD. The variability associated with all other morphologies indicates that when LDs are larger than 200 nm, deformations do not depend on the sizes of interacting LDs (diameter ratio ‘small LDs indenting large LD’ vs. any other morphology P< 0.0001; all other comparisons P> 0.4).

When we measured the distances between the monolayers, we found that the monolayers were closer for untagged Cidec than for Cidec-EGFP (mean of distances untagged Cidec 8.5 nm, SD 2.55 nm, n=8, P=0.0164 compared to Cidec-EGFP) (Fig. 2i). The observed difference compared to Cidec-EGFP interfaces is in good agreement with the dimensions of the GFP molecules (16), suggesting that the increase in distance is due to the space taken up by EGFP moieties. These results suggest that the spacing between LD monolayers is determined by dense packing of Cidec molecules, and is influenced by the size of the Cidec construct mediating the interaction.

Having identified determinants of the architecture of the LD interface, we next sought to link them to neutral lipid transfer function. We hypothesised that if Cidec-EGFP has an effect due to the bulky size of the tag (27 kDa) as compared to untagged Cidec, Cidec conjugated to SUMOstar may have a reduced effect due to the intermediate size of the tag (12 kDa). By titrating Doxycycline dosage, comparable *Cidec, Cidec-SUMOstar* and *Cidec-EGFP* transcript levels were confirmed for all three stable cell lines (Suppl. Fig. S3). For FM imaging, we fixed cells expressing untagged Cidec and Cidec-SUMOstar 24h after oleic acid and Doxycycline were added. Similarly to Cidec-EGFP (Fig. 1b), for both Cidec-SUMOstar and Cidec we observed upon Doxycycline addition an increase in the mean LD volume (n=3 repeats; P< 0.0001 for Cidec-SUMOstar, P=0.0620 Cidec)(Fig. 3a) and a reduction in the mean number of LDs per cell (n=3 repeats, P=0.0026 for Cidec-EGFP, P=0.0005 for Cidec-SUMOstar, P=0.0017 for Cidec) (Fig. 3b). No difference in mean LD volume (Fig. 3a)(n=3, P=0.4982) or mean LD number (n=3 repeats; P=0.9306)(Fig. 3b) was observed upon induction of cytosolic EGFP expression. When comparing cells expressing the different constructs, we found that the LD volumes (n=3 repeats; P=0.0009 for group comparison; means EGFP: 0.37 *µ*m^3^, Cidec-EGFP: 1.7 *µ*m^3^, Cidec-SUMOstar: 2.2 *µ*m^3^, Cidec: 3.5 *µ*m^3^)(Fig. 3a) as well as the number of LDs differed (n=3 repeats; P=0.022 for group comparison; means EGFP: 139.8 LDs/cell, Cidec-EGFP: 51.1 LDs/cell, Cidec-SUMOstar: 29.4 LDs/cell, Cidec: 23.6 LDs/cell) (Fig. 3b). The number of LDs per cell was lower in cells expressing Cidec-SUMOstar as compared to Cidec-EGFP, indicating a potential influence of the tag size (P=0.0329) (Fig. 3b). These results confirm that all three constructs enlarge LDs and reduce their number per cell, albeit with varying efficiency.

**Figure 3:**
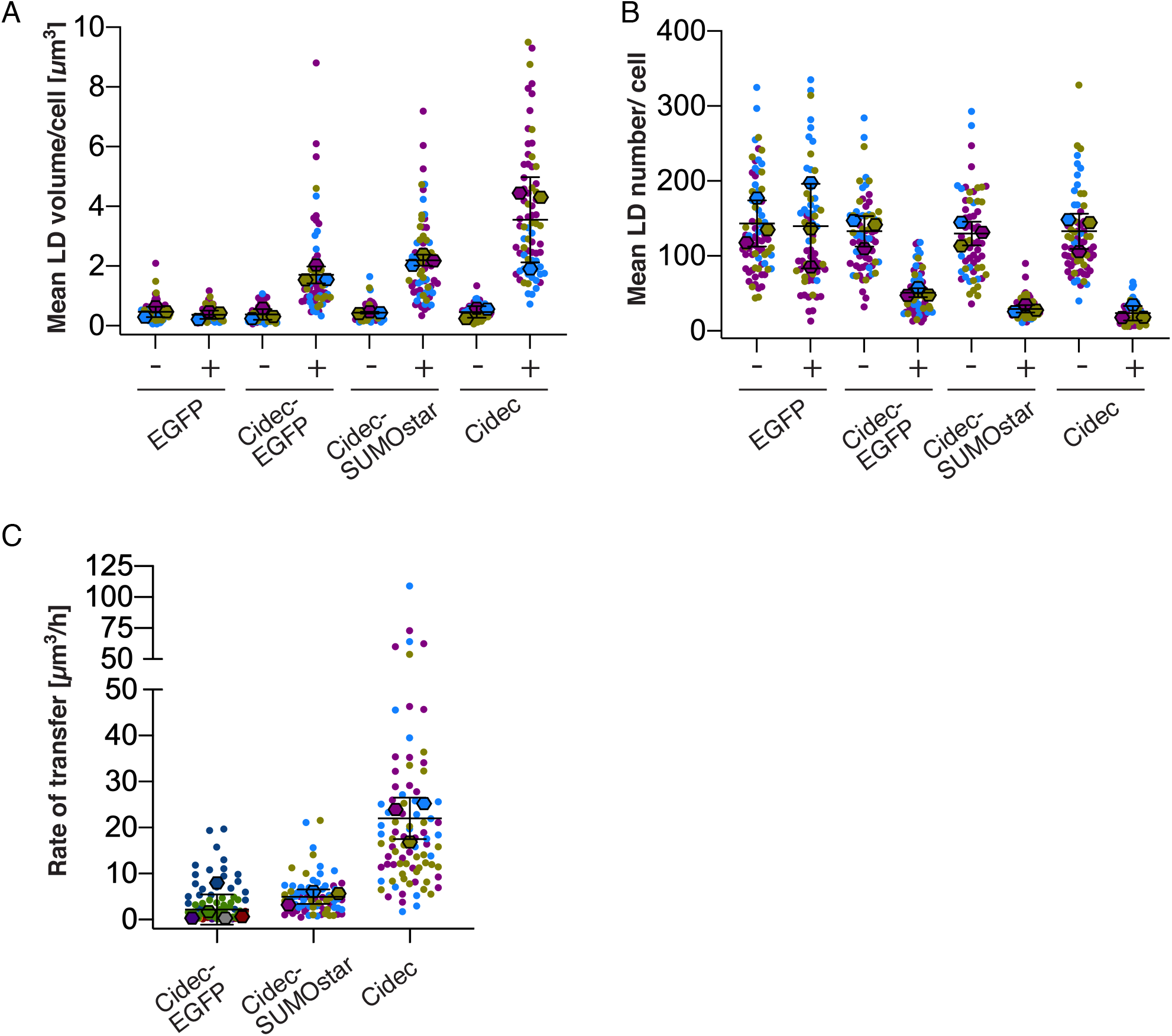
Quantitative fluorescence microscopy of HeLa cells expressing Cidec-EGFP, Cidec-SUMOstar or Cidec. **(A and B)** Quantification of FM images of fixed HeLa cells, loaded with oleic acid and induced with Doxycycline for 24 hours prior to fixing and imaging. Means of individual experimental repeats are represented by diamond symbols, individual cells are represented by dots, colour coded according to experiment. In each experimental repeat, 11-35 cells were analysed per condition. **A)** The mean LD volume per cell upon expression (+Dox) or no expression (-Dox). Black bars represent the mean of three experimental repeats with standard deviation (SD) (EGFP: -Dox 0.47 *µ*m^3^, SD 0.17 *µ*m^3^, +Dox 0.37 *µ*m^3^, SD 0.14 *µ*m^3^; Cidec-EGFP: -Dox 0.38 *µ*m^3^, SD 0.18 *µ*m^3^, +Dox 1.71 *µ*m^3^, SD 0.28 *µ*m^3^; Cidec-SUMOstar: -Dox 0.44 *µ*m^3^, SD 0.03 *µ*m^3^ +Dox 2.20 *µ*m^3^, SD 0.18 *µ*m^3^; Cidec: -Dox 0.46 *µ*m^3^, SD 0.19 *µ*m^3^, +Dox 3.55 *µ*m^3^, SD 1.43 *µ*m^3^). Cidec-EGFP data is also shown in Fig 1B. **B)** The mean LD number per cell upon expression (+Dox) or no expression (-Dox). Black bars represent means of three experimental repeats with standard deviation (SD) (EGFP: -Dox 143.2 LDs/cell, SD 30.7 LDs/cell, +Dox 139.8 LDs/cell, SD 56.3 LDs/cell; Cidec-EGFP: -Dox 132.9 LDs/cell, SD 20.3 LDs/cell, +Dox 51.1 LDs/cell, SD 5.8 LDs/cell; Cidec-SUMOstar: -Dox 130.0 LDs/cell, SD 15.7 LDs/cell, +Dox 29.4 LDs/cell, SD 4.9 LDs/cell, Cidec: -Dox 132.9 LDs/cell, SD 23.5 LDs/cell, +Dox 23.6 LDs/cell, SD 9.7 LDs/cell). **C)** The rate of lipid transfer from donors to acceptors, as a change of donor volume. Measurements done in live cells imaged by time-lapse FM of LipidTOX signals (Movies 8-10). Live FM started 2h post-Doxycycline induction for Cidec and Cidec-SUMOstar, and 24h post-Doxycycline induction for Cidec-EGFP. Mean of individual experimental repeats are represented by diamond symbols, individual lipid transfer events are represented by dots, colour coded according to experiment. Black bars represent mean of all experimental repeats and SD (Cidec-EGFP: 2.17 *µ*m^3^/h, SD 3.29 *µ*m^3^/h, n=5 repeats each analysing 4-43 cells; Cidec-SUMOstar: 4.95 *µ*m^3^/h, SD 1.56 *µ*m^3^/h, n=3 repeats each analysing 18-36 cells; Cidec: mean 21.98 *µ*m^3^/h, SD 4.52 *µ*m^3^/h, n=3 repeats each analysing 23-34 cells). Only donors involved in active lipid transport and in contact with a single acceptor were considered.

We next asked whether the varying efficiency was due to LD merging occurring at different speeds in HeLa cells expressing Cidec-EGFP, Cidec-SUMOstar or Cidec. We thus aimed to determine the rate of neutral lipid transfer mediated by the different constructs. In 3T3-L1 pre-adipocytes overexpressing Cidec-EGFP, lipid exchange rates of 0.13 *µ*m^3^/s were reported (3), whereas in differentiated 3T3-L1 adipocytes expressing endogenous Cidec, lipid exchange rates of 5.6 *µ*m^3^/s have been reported (8). In pre-adipocytes this lipid exchange resulted in net neutral lipid transfer to the larger droplet at a rate of 4.8 *µ*m^3^/h (3). We collected live cell FM images of LDs, starting 2h after induction of expression of Cidec and Cidec-SUMOstar. For Cidec-EGFP, LDs were very small at this time point, so we started live FM 24h after induction. We identified LD pairs engaged in active lipid transfer by the close proximity of LD cores labelled with LipidTOX (Movies 8-10). All three Cidec constructs mediated lipid transfer in a net directional manner from the smaller LDs (donors) to the larger LDs (acceptors), as expected from previous reports (2, 3). We measured the change of donor volume over time (see Materials and Methods). Importantly, only donors that were in contact with a single acceptor were taken into account. For Cidec-EGFP, we found the mean volume reduction of the donor to be 2.2 *µ*m^3^/h (n=5 repeats) (Fig. 3c), indicating that a complete transfer of neutral lipid content from donors to acceptors occurred in the range of hours. For Cidec-SUMOstar, the mean volume reduction rate was 5.0 *µ*m^3^/h (n=3 repeats), while in the presence of untagged Cidec, the mean volume reduction rate was 22.0 *µ*m^3^/h (n=3 repeats) (Fig. 3c). Such a rate indicates that untagged Cidec transfers neutral lipids between donor and acceptor LDs within minutes. The rates thus differed significantly (P=0.0068).Taken together, these data suggest that the rate of net lipid transfer slows down considerably with increased bulkiness of the protein at LD-LD interfaces.

We next investigated the Cidec-mediated change in LD volume over time by analysing the kinetics of the process. To this end we implemented a semi-automated image analysis pipeline (see Materials and Methods), which we applied to the live FM data obtained for untagged Cidec and Cidec-SUMOstar. Due to the relatively slow merging of LDs in Cidec-EGFP cells and the resulting use of different imaging settings, we did not include Cidec-EGFP. Our analysis automatically determined the time points when a pair of LDs engaging in active lipid transfer first came into contact and when the lipid transfer event was completed (Fig. 4a and b). For pairs identified as engaged in transfer, we obtained LD volumes in each movie frame and plotted them against time. The resulting curves showed that the change in LD volume of both donors and acceptors was not linear over time, but accelerated over the course of the transfer event. This was the case both for Cidec (Fig. 4c and d) and Cidec-SUMOstar (Fig. 4e). We fitted an exponential function to the curves and calculated rate constants from the fit (R values) (see Materials and Methods). The smaller the R value, the slower the whole transfer event. Within an LD pair engaged in transfer, the R value of the donor should correspond to the R value of the acceptor, unless the LDs are engaged in additional transfer events. Hence, to determine whether LDs are engaged in a single transfer event at a time, for each event we plotted logarithmically the R values of donor vs. acceptor (Fig. 5a). For the majority of events, donor and acceptor R values were very similar. To exclude events likely involved in multiple transfers, we further considered only those transfer events for which log10(R_Acceptor_)/log10(R_Donor_)-1 was smaller than 0.4 (Fig. 5a, filled circles). Among these, the mean R_Donor_ value for Cidec was 0.030 /s, and 0.007 /s for Cidec-SUMOstar (Fig. 5b). R_Donor_ did not correlate with the ratio between the starting volumes of the acceptor and the donor (Fig 5b), indicating that the initial size difference is not the dominant determinant of overall transfer speed. We next determined the instant change of volume for each donor LD at defined time points corresponding to temporal fractions of the transfer event. In this way, we aimed to normalise for the variable duration of transfer events. To further account for the diversity of donor sizes at the start of the event, we divided the instant change of volume by the donor volume at that same time point. We then plotted these values against the progression of the event, represented as percentage of the time between first contact and completion. The resulting curves allow comparing the change of volume between different events by normalising for the absolute donor volume and for the time point during the event (Fig. 5c and d, grey lines). From these data, we calculated the median behaviour of donor volumes in the presence of Cidec (Fig. 5c, red line) and Cidec-SUMOstar (Fig. 5d, red line). These data show that there is inherent variability in the volume change among events, which is not accounted for by the absolute size of the donor, nor by the relative time point during the event. The distributions of curves for Cidec and Cidec-SUMOstar overlap partly, indicating that individual events can be indistinguishable. Collectively, however, Cidec events show a faster relative change of volume than Cidec-SUMOstar events, at all time points assessed (Fig. 5e, P< 0.0001 at all points compared).

**Figure 4:**
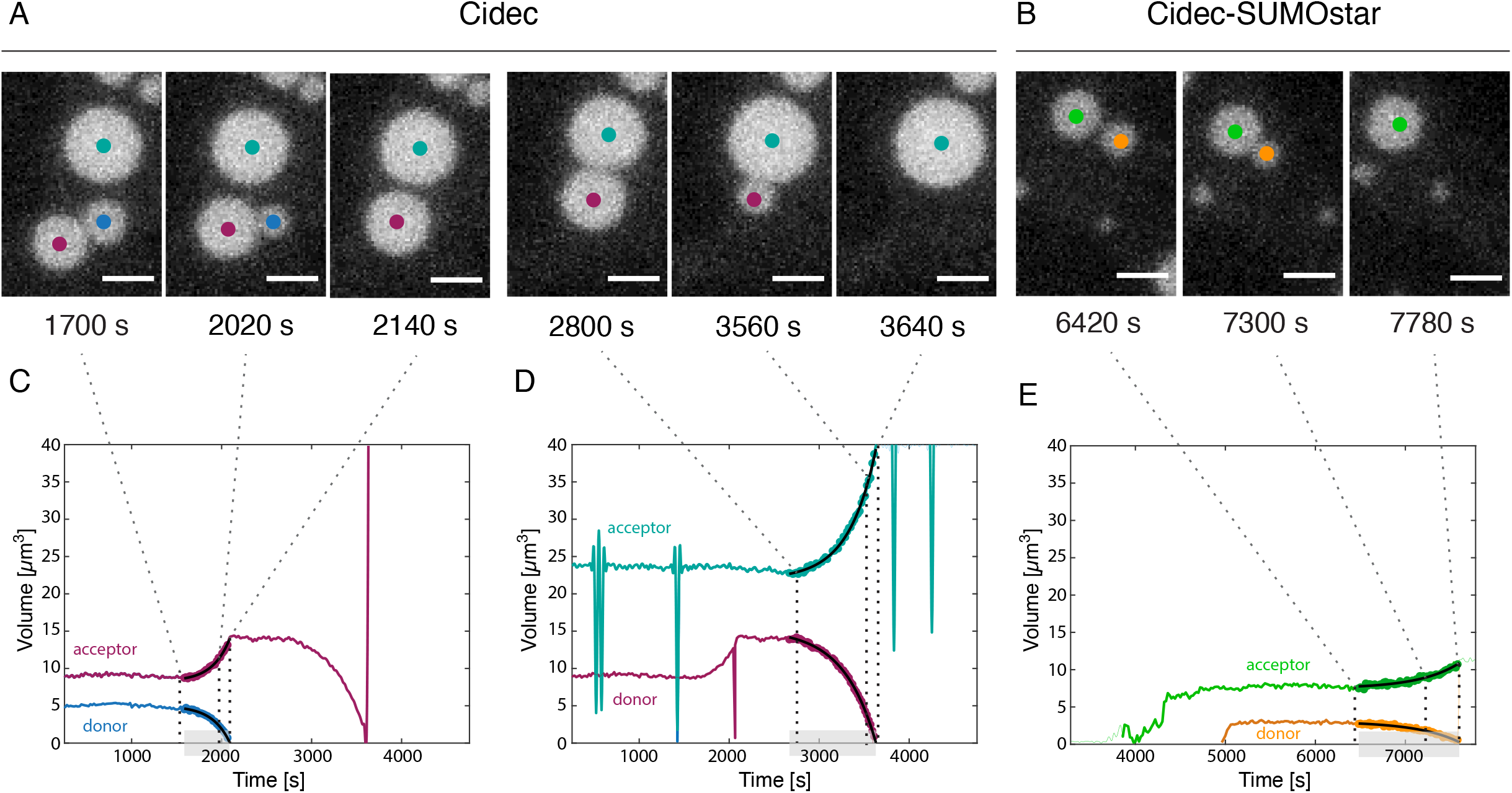
Lipid transfer in HeLa cells expressing Cidec or Cidec-SUMOstar follows exponential kinetics. **A) and B)** Time-course live FM of HeLa cells expressing untagged Cidec (A) and Cidec-SUMOstar (B). Representative LD pairs engaged in active lipid transfer. Panels are maximum projections of a z-stack. Scale bars: 2.5 *µ*m. **C), D) and E)** Traces of volume change of individual LDs over time, in Cidec (C, D) and Cidec-SUMOstar (E) expressing cells. Grey overlays on x-axes highlight the time when LDs are in contact (“event time”), and involved in active transfer. Black lines overlaid on traces correspond to the exponential fit, which is used to determine R-values (shown in Fig 5A and B).

**Figure 5:**
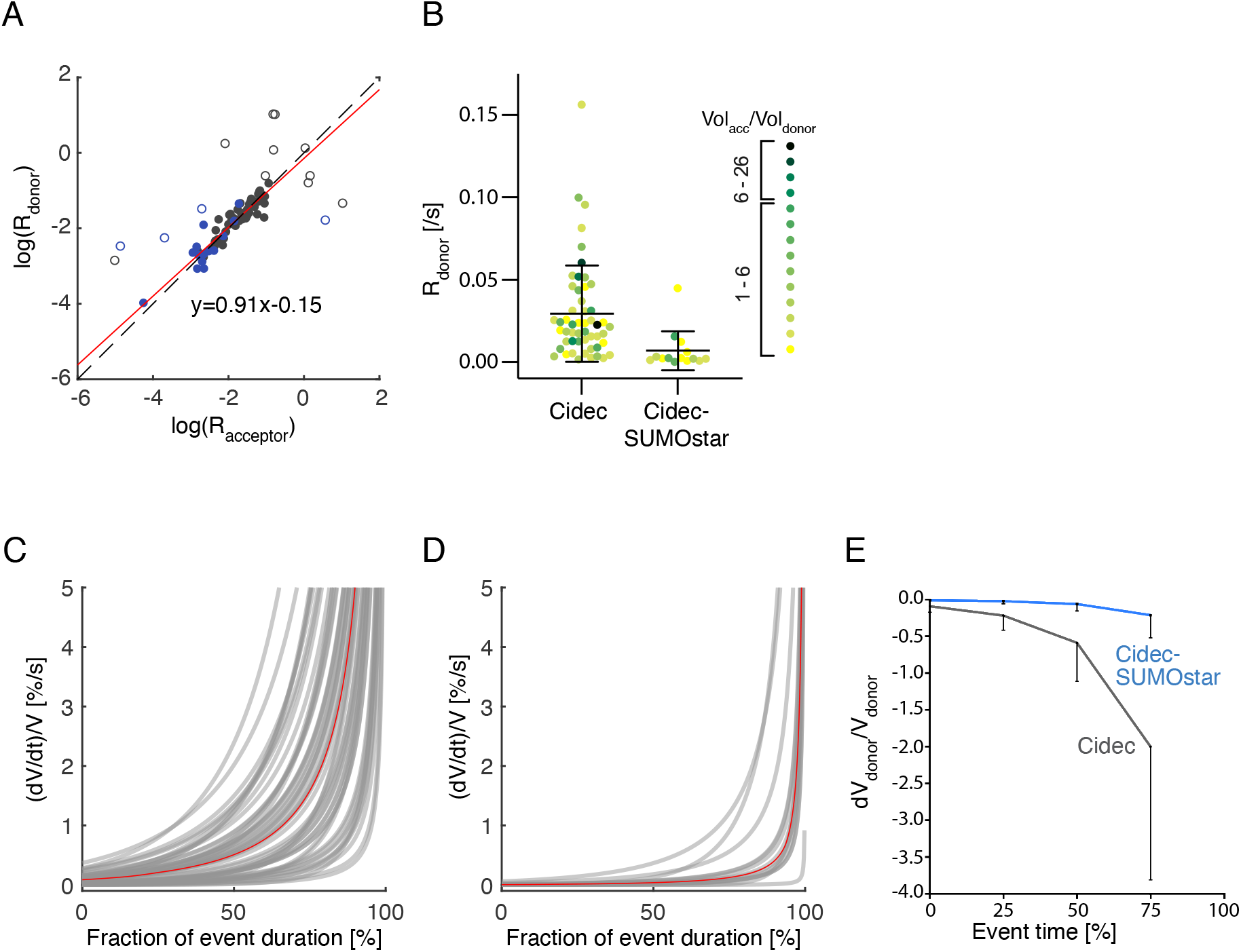
Quantitative analysis of lipid transfer kinetics in HeLa cells expressing Cidec or Cidec-SUMOstar. **A)** Detection of outliers. Log10 of R-values, calculated from exponential fits as shown in Figure 4C, D and E, of acceptor LD are plotted against log10 of R-values of the corresponding donor LD. Cidec: black circles. Cidec-SUMOstar: blue circles. Pairs differing by more than 0.4 (empty circles) were excluded from further analysis (for details see text and materials and methods). **B)** R-values for Cidec (n=52) and Cidec-SUMOstar (n=14). Each data point corresponds to one event, and is color-coded according to the starting ratio of acceptor volume to donor volume. Black bars correspond to mean and standard deviation (Cidec: n=52; mean: 0.007 /s, Cidec-SUMOstar: n=14; mean: 0.030 /s. P=0.0064). **C) and D)** Relative rate of volume change vs time for individual donors: Cidec (C) and Cidec-SUMOstar (D). Individual curves for each donor plotted as change of volume over time of event duration. Volume is expressed as the change of volume divided by the final volume of the merged LDs. Time is given as a fraction of the event at a particular volume change. Red curves represent an median curve calculated from all individual donor curves (C: n=52, D: n=14). **E)** Mean instant change of volume (relative to the current donor volume) calculated at fractions (0-75%) of the event duration (event time). Black bars represent standard deviation.

Taken together, these results show that the transfer of lipids at LD-LD interfaces follows exponential kinetics, starting slowly and accelerating towards the end. While this behaviour is robust and retained in the presence of protein tags, the magnitude of the transfer rate is altered throughout the process by the bulkiness of the protein at the interface. These findings are in line with our results from cryo-ET, which show that monolayer integrity and overall architecture are consistent features of LD interfaces, while the magnitude of the gap between the monolayers is influenced by the bulkiness of the protein at the interface. Furthermore, variability between individual events is not attributed to the starting absolute volume or volume ratio, suggesting that overall speed is influenced by additional factors. One such factor could be interface morphology, which we found by cryo-ET to also be independent of the volume ratio.

The contents of LD cores are among the most hydrophobic molecules found in cells. Thus how “merging” of LDs in the aqueous cytosol is controlled poses a unique cell biological problem. Here we show that transfer of neutral lipids occurs through two largely intact monolayers, rather than a pore akin to membrane fusion. This observation does not rule out the occurrence of molecular-level disturbances in the monolayers, which could cause leakiness and thereby transit of TAG molecules. By the close apposition of monolayers, LD interfaces show resemblance to membrane contact sites (3). However, the distances between the monolayers are considerably smaller than those typically reported for membrane contact sites involved in lipid transfer (17). Furthermore, in contrast to lipid transfer proteins at membrane contact sites (18), Cidec is not known to have a lipid transfer domain. Hence the transfer of TAGs is likely driven by a fundamentally different mechanism from other lipid transfer events. Dense packing of Cidec between and within the two monolayers possibly creates a highly hydrophobic microenvironment that allows flux of hydrophobic molecules. The intactness of the monolayers might be key for retaining a difference in surface tension between the two LDs. This difference generates Laplace pressure, which has been proposed as the driving force for the transfer (3). Our finding that transfer follows exponential kinetics provides prerequisite evidence for this model (11). In line with this model, our observation that LDs smaller than 200 nm do not deform in shape but impose their curvature on larger LDs indicates that small LDs have a greater internal pressure or surface tension than large LDs. The interface architecture and transfer kinetics are quantitatively altered by changes in Cidec size through addition of tags, but qualitatively remain unperturbed. These findings further suggest a transfer mechanism intrinsic to the physical properties of LDs and of the interface formed by Cidec.

In summary, our results suggest that the transfer of TAGs through the LD monolayers is driven by pressure and enabled by a specialised microenvironment formed by Cidec between them. Thus Cidec may be facilitating a form of Ostwald ripening as the mechanism underlying the unilocularity of white adipocytes.

## Acknowledgements

We would like to thank the MRC LMB electron microscopy facility for support with cryo-ET data acquision. Work in the group of W.K. was supported by the Medical Research Council (MC_UP_1201/8). W.K. is at present supported by the University of Bern. D.B.S (WT 107064) is supported by the Wellcome Trust, the MRC Metabolic Disease Unit [MRC_MC_UU_12012.1], and The National Institute for Health Research (NIHR) Cambridge Biomedical Research Centre and NIHR Rare Disease Translational Research Collaboration. The IMS MRL Imaging Core is supported by a Wellcome Trust Major Award [208363/Z/17/Z].

## Materials and Methods

### Cloning

All Cidec constructs used in this study contain the *Mus musculus* Cidec sequence. Mouse Cidec cDNA was amplified by PCR using Phusion High-Fidelity DNA Polymerase (Thermo Fisher, F530L). The untagged Cidec transcript flanked by SacII and NotI restriction sites at the amino- and carboxyl-terminus, respectively, was cloned into Gateway Entry vector, pEN-Tmcs, using a DNA Ligation Kit, Mighty Mix (Takara Bio, 6023) and transformed into One Shot TOP10 Chemically Competent *E*.*coli* (Thermo Fisher, C404003). For EGFP tagging at the carboxyl-terminus, a SacII and BamHI restriction site flanked Cidec sequence was inserted into a pEGFPN3 vector. The sequence-verified insert was subcloned into a pEN-Tmcs vector using SacII and NotI restriction sites. In order to generate expression clones, sequence-verified inserts along with Tet-responsive elements flanked by *att*L sites in the Gateway Entry vector were recombined into a Gateway Destination vector, pSLIK-hygromycin containing *att*R sites using Gateway LR Clonase II enzyme mix (Thermo Fisher, 11791020). Recombination reactions were transformed into One Shot Stbl3 Chemically Competent *E*.*coli* (Thermo Fisher, C737303).

### Generation of *Cidec* null MEF and HeLa cell lines with Doxycycline-inducible expression of Cidec constructs

*Cidec* null MEFs and HeLa cell lines with Doxycycline-inducible expression of Cidec (untagged, SUMOstar- and EGFP-tagged Cidec) were generated by lentiviral transduction. To generate lentiviruses, HEK293T cells at approximately 70% confluency were transfected using Lipofectamine LTX with Plus Reagent (Thermo Fisher, 15338100) according to the manufacturer’s protocol. A typical transfection reaction included 7.5 *µ*g pMDLg/pRRE, 7.5 *µ*g pRSV-Rev, 5 *µ*g pVSV-G, 1 *µ*g pEGFP and 10 *µ*g pSLIK-hygromycin plasmid DNA (with integrated untagged, SUMOstar- and EGFP-tagged Cidec WT cDNA sequences). 24 hours post-transfection, the culture medium was removed and the cells were replenished with UltraCULTURE serum-free cell culture medium (Lonza, BE12-725F). Medium containing secreted lentivirus was collected every 24 hours for a total of 72 hours and stored at 4°C. Lentivirus-containing media was centrifuged at 2000 g at 4°C for 20 minutes and the supernatant was filtered through 0.45 *µ*m syringe filters.

To concentrate the lentiviral supernatant, Centricon Plus-70 Centrifugal Filters (Merck Millipore, UFC710008) were used according to the manufacturer’s protocol. Sample filter cups were pre-rinsed with PBS and lentiviral supernatant was loaded onto the sample filter cups, sealed and centrifuged at 3500 g for 30 minutes at 15°C. To recover the concentrated lentivirus, the concentrate collection cups were inverted and placed on top of the sample filter cups, and then centrifuged at 3500 g for 10 minutes at 15°C. The concentrated lentivirus was either used to transduce *Cidec* null MEFs and HeLa cells at 2 to 3 different viral titres, or aliquoted into cryovials for long term storage at - 80°C. Cells were selected 24 hours post-lentiviral transduction using 200 *µ*g/mL of hygromycin. Non-lentivirus transduced cells were used as a control for antibiotic selection.

### Cell culture

*Cidec* null MEFs were a kind gift from Masato Kasuga. They were grown at 37°C and 5% CO_2_ in high glucose DMEM media containing 10% Tetracycline-free FBS (Pan Biotech, P30-3602), 1 mM sodium pyruvate, 2 mM L-glutamine, 1x non-essential amino acids and 50 *µ*M β-mercaptoethanol. The primary MEFs were immortalised by transfection of 2 μg of simian virus 40 large T-antigen-expressing vector employing Fugene 6 Transfection Reagent (Promega, E2311) followed by five rounds of a 1 in10 split to achieve 1/100,000-fold splitting. Untransfected primary MEFs that underwent the 1/100,000-fold splitting were used as a control to ensure that all surviving cells were immortalised. Immortalised *Cidec* null MEFs were then transduced with pBABE-mPPARγ2 to ensure they had the potency to differentiate into adipocytes. BOSC 23 retroviral packaging cells were ∼70% confluent when transfected with 12 *µ*g of pBABE-mPPARγ2 or pBABE-EGFP plasmid DNA using Fugene 6 transfection reagent (Promega, E2311). Media containing secreted retrovirus was collected 72 hours post-transfection and filtered through 0.45 *µ*m syringe filters. The filtered retroviral stocks were used to transduce ∼50-60% confluent immortalised *Cidec* null MEFs with the addition of 12 *µ*g/mL polybrene. Puromycin selection was initiated 24 hours post-transduction at a concentration of 4 *µ*g/mL.

To induce differentiation of *Cidec* null MEFs into adipocytes, cells were grown to 2-days post-confluence, then induced to differentiate in culture medium supplemented with 8 *µ*g/mL biotin (Merck, B4639), 8 *µ*g/mL D-pantothenic acid (Merck, P5155), 1 *µ*M rosiglitazone (Merck, R2408), 0.5 mM 3-isobutyl-1-methylxanthine (Merck, I5879), 1 μM dexamethasone (Merck, D4902) and 1 *µ*M insulin. Two days thereafter, culture medium was changed to contain 8 *µ*g/mL biotin, 8 *µ*g/mL D-pantothenic acid, 1 *µ*M rosiglitazone and 1 *µ*M insulin. From day 4 of differentiation onwards, culture medium was changed every 2 days until the cells were used for experiments.

HeLa Doxycycline-inducible stable cell lines were grown as an adherent culture at 37°C, 5% CO_2_ in a high glucose DMEM media containing pyruvate, GlutaMAX (ThermoFisher 31996). The media was additionally supplemented with 10% Tet-approved heat-inactivated FBS (Pan Biotech p30-3602), 10 mM HEPES pH 7.2, 0.2 mg/ml hygromycin B (Invitrogen 10687010) and 1x non-essential amino acids (ThermoFisher 11140). Cell lines were regularly tested for mycoplasma infection using the MycoAlert mycoplasma detection kit (Lonza, LT07-418).

### Determination of mRNA transcript levels of *Cidec* constructs in HeLa cells

In order to determine and achieve comparable mRNA expression levels of *Cidec* in untagged, SUMOstar- and EGFP-tagged HeLa cell lines, Doxycycline of 0.5, 1.0 or 2.0 *µ*g/mL was added to seeded cells in the presence of 200 *µ*M of oleic acid. 24 hours post-induction, RNA was harvested using an RNeasy Mini Kit (QIAgen, 74106) by following the manufacturer’s protocol. 400 ng of RNA was treated with 1 unit of RQ1 RNase-Free DNase (Promega, M610A) at 37°C for 30 minutes and was inactivated by RQ1 DNase Stop Solution at 65°C for 10 minutes. To generate cDNA standards and a negative control for reverse transcription, 1 *µ*g of pooled RNA was prepared. RNA was reverse transcribed into cDNA using a LunaScript RT SuperMix Kit (NEB, E3010L) by following the manufacturer’s protocol, using thermocycling conditions as follows: 25°C for 2 minutes, 55°C for 10 minutes and 95°C for 1 minute. cDNA was diluted 10x and qPCR was performed on an Applied Biosystem QuantStudio 7 Flex Real-Time PCR System. A typical TaqMan qPCR reaction included 1x TaqMan Universal PCR Master Mix (Thermo Fisher, 4304437) and 1x TaqMan Gene Expression Assay, in which *GAPDH* was used as a housekeeping gene (Cidec: Mm01184685_g1; GAPDH: Hs02758991_g1). mRNA levels were normalized to the levels in untagged *Cidec* line untreated with Doxycycline. From this, it was determined that 0.5 *µ*g/mL of Doxycycline in untagged Cidec and Cidec-EGFP lines induced comparable mRNA expression levels of *Cidec* as 2.0 *µ*g/mL of Doxycycline in Cidec-SUMOstar line. Thus, these Doxycycline concentrations were used for the described fixed and live cell imaging experiments.

### Fixed cell imaging

Cells were seeded onto ethanol-treated glass coverslips (Thermo Fisher) in 12-well plates. *Cidec* null MEFs were differentiated into mature adipocytes according to the protocol above. 1*µ*g/mL of Doxycycline was either added or omitted throughout the course of differentiation. An hour before fixation, LDs were stained with 1x HCS LipidTOX Deep Red Neutral Lipid Stain (Thermo Fisher, H34477) at 37°C. To study the effects of Cidec on LD enlargement at a fixed time point in Hela cells expressing Doxycycline-inducible Cidec constructs, the day after seeding, cells were washed twice with PBS and loaded with 200 μM oleic acid (Merck, O3008) and various Doxycycline concentrations (0.5-2 *µ*g/mL) in order to achieve comparable Cidec transcript levels (see above). 23 hours post-Doxycycline induction, LDs were stained with 1x HCS LipidTOX Deep Red Neutral Lipid Stain for an hour at 37°C.

Cells were fixed with 4% (v/v) formaldehyde (Merck, 47608) diluted in PBS for 15 minutes at room temperature. Cells were then washed 3 times for 5 minutes with PBS and mounted using VECTASHIELD Antifade Mounting Medium (Vector Laboratories, H-1000) or Prolong Gold Antifade Mountant with DAPI (Thermo Fisher, P36931). For *Cidec* null MEF-derived adipocytes, 2-dimensional images were acquired using a Leica SP8 confocal microscope. EGFP and LipidTOX Deep Red Neutral Lipid dyes were excited at 488 and 637 nm, and emission signals were collected at 495-535 and 645-700 nm, respectively. For HeLa cell lines, 3-dimensional images were acquired using a Leica SP8 confocal microscope at 0.3-0.4 *µ*m z-sections. EGFP and LipidTOX Deep Red Neutral Lipid dyes were excited at 488 nm and 637 nm, and emission signals were collected at 495-550 nm and 645-690 nm, respectively. Experiments were repeated twice for *Cidec* null MEFs-derived adipocytes with 25 cells being analysed for LD volume in each experiment and three times for HeLa cells with at least 11 cells being analysed in each experiment.

### Analysis of LD sizes and number in fixed cell fluorescence images

In 2-dimensional images of fixed *Cidec* null MEFs-derived adipocytes, LD sizes were derived by measuring LD diameters using Fiji software (19) based on LipidTOX Deep Red dye staining. A line was drawn across the LDs on focal planes and these measurements were used to calculate LD volumes by assuming that LDs are spherical in shape. In images of fixed HeLa cell lines, LD sizes and numbers were determined by using Bitplane Imaris software version 9.6.0 (Oxford Instruments), based on LipidTOX Deep Red dye staining. The 3-dimensional images were segmented by ‘Spots’ creation wizard with ‘Different Spot Sizes (Region Growing)’ algorithm setting enabled. ‘Estimated XY Diameter’ was set between 0.6 and 2.0 *µ*m with ‘Background Subtraction’ enabled. Images were further filtered using ‘Quality’ filter type of at least 2.7 and ‘Spot Region Type’ of ‘Absolute Intensity’. Diameters of the ‘Spot Regions’ were determined from ‘Region Border’ at thresholds of 16-35. LD numbers per cell were generated based on the number of spots detected.

### Live cell imaging

Cells were seeded onto 35 mm imaging dishes with a polymer coverslip bottom (Ibidi, 81156). 24 hours before live cell imaging was performed, 200 μM oleic acid was added to cells in culture medium supplemented with 10% Tetracycline-free FBS (Pan Biotech, P30-3602), 2 mM L-glutamine (Merck, G7513), 100 units/mL penicillin and 0.1 mg/mL streptomycin (Merck, P4333) and 200 *µ*g/mL Hygromycin B (Merck Millipore, 400052). Two hours before imaging of HeLa cells with Doxycycline-inducible expression of untagged Cidec or Cidec-SUMOstar, 0.5 or 2 *µ*g/mL of Doxycycline, respectively, was added to the cells in the presence of 1x HCS LipidTOX Deep Red Neutral Lipid Stain. In the case of Cidec-EGFP, 0.5 *µ*g/mL of Doxycycline was added 24 hours before the imaging. Live cell imaging was performed on a Leica SP8 confocal microscope in a chamber maintained at 37°C with 5% CO_2_, using a 63x objective with NA=1.4, and using an additional two-fold or three-fold zoom on the microscope level. LipidTOX Deep Red Neutral Lipid Stain was excited at 633 nm, and the emission signal was collected 640-700 nm. For HeLa cells expressing untagged Cidec or Cidec-SUMOstar, multiple positions of 3-dimensional images were acquired at 20 second intervals at 0.4 *µ*m z-sections for at least 2 hours, while for Cidec-EGFP, 3-dimensional images were acquired at 2.5-minute intervals for at least 20 hours. During all acquisitions of untagged Cidec and Cidec-SUMOstar data, a three-fold zoom was used, resulting in a pixel size of 120 nm, while in 3 out of 4 data acquisition sessions for Cidec-EGFP, a two-fold zoom on the microscope was used, resulting in a pixel size of 180 nm, as compared to 120 nm for all other data.

### Manual analysis of lipid transfer rate

To determine the rate of neutral lipid transfer, maximum projection images were generated from the 3D stacks using Fiji (19) and the diameters of donor droplets were measured at least at two different positions using Fiji. The average of the two or more measured diameters were used to derive the volume of the lipid donors. The change of the lipid donor volume over time (Δvolume/Δtime) was calculated by assuming that LDs are spherical in shape. In all cases, the rate of neutral lipid transfer was determined for donors that underwent active lipid transfer and were tethered to a single acceptor. In the case of Cidec-EGFP, only LD pairs with enriched Cidec-EGFP at the LD interfaces were analysed.

### Semi-automated analysis of lipid transfer rate

Kinetics of neutral lipid transfer were analysed with a custom-made MATLAB (MathWorks) script. Movies were acquired as described above and individual LDs were further segmented out and tracked over the course of the 3D image sequence. An event at time *t*_*max*_ is defined as a merging of two tracks into a single one, and the donor droplet is identified as the droplet whose volume will vanish at *t*_*max*_. The evolution of the volume of the donor droplet over time is modelled as 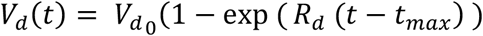 where 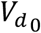 is the initial volume of the donor and *R*_*d*_ is the transfer rate. Similarly, the volume of the acceptor droplet is modelled as 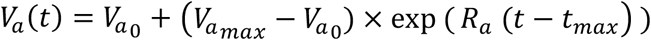where 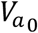 and 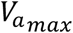 are respectively the initial and final volume of the acceptor and *R*_*a*_ is the transfer rate. In both cases, a least-square procedure allows to retrieve the 5 parameters for each event. Each LD pair is considered as an individual event. To exclude outliers, corresponding to LDs in contact with more than one other LD, the rate *R*_*d*_ for the donor was plotted against the rate *R*_*a*_ for the acceptor for each event. We considered only those transfer events for which | log_10_*R*_*a*_*/*log_10_*R*_*d*_−1| <0.4, thereby excluding events likely to be involved in multiple transfers. The two rates of all events were plotted in Prism as scatter plots with error bars displaying mean and standard deviation. The rates obtained for Cidec untagged and Cidec-SUMOstar were compared using an unpaired two-tailed Mann Whitney test (p-value < 0.0001).

Furthermore, the instant change of volume of the donor at certain time points was determined. From the model parameter, we estimated the temporal derivatives of the evolution volume of the donor dVd/dt and normalized it by the current volume Vd in order to compute a relative change of volume over time denoted dVd/Vd. In order to synchronize the estimated evolution of volume across events, we arbitrarily defined a reference start time for each event as the time when the donor is at 95% its initial volume. Synchronized instant relative change of volume were then represented together and a median curve was computed for each condition. Specific relative change of volume was estimated at 25%, 50%, 75% and 100% of the time relative to this reference starting time.

### Sample preparation for cryo-fluorescence microscopy and cryoFIB milling

HeLa Doxycycline-inducible stable cell lines were grown for approximately 48 h on holey carbon gold EM grids (200 mesh, R2/2, Quantifoil) prior to plunge freezing. 24 h after seeding, HeLa cells expressing Cidec-EGFP were fed with 0.4 mM oleic acid (Sigma, OA O3008) and induced with 1 *µ*g/mL Doxycycline for Cidec-EGFP expression. 16 h post-induction, these cells were stained for 1 h with LipidTOX Deep Red dye (ThermoFisher scientific, H34477) and plunge frozen. HeLa cells expressing Cidec untagged were grown for 24 h and then fed with 0.4 mM oleic acid. 24 h later, these cells were induced with 1 *µ*g/mL Doxycycline for Cidec untagged expression. 30min post-induction, cells were stained for 1 h with LipidTOX Deep Red dye and 1h 30min post-induction, cells were plunge frozen. Plunge freezing was performed with custom-made manual plunger and cryostat (20). To this end, grids were manually back side blotted with Whatman filter paper No 1 and immediately vitrified in liquid ethane. Grids were screened at −195°C for ice quality and for regions featuring cells of interest by cryo-fluorescence microscopy (cryoFM) in a Leica EM cryo-CLEM system (Leica Microsystems) in a humidity-controlled room. The system was equipped with a HCX PL APO 50x cryo objective with 0.9 NA (Leica Microsystems), an Orca Flash 4.0 V2 SCMOS camera (Hamamatsu Photonics), a Sola Light Engine (Lumencor), a L5 filter (Leica Microsystems) for the detection of EGFP and a Y5 filter (Leica Microsystems) for far red detection. 1.8 x 1.8 mm montages of the central part of the grids were taken. These montages were later manually correlated with scanning electron beam micrographs and served for finding the regions of interest for lamella preparation by cryo-FIB milling. Z-stacks in 1 *µ*m steps were acquired of regions of interest corresponding to cells with enlarged LDs and Cidec-EGFP accumulation at LD-LD interfaces.

### Cryo-FIB milling

Thin lamellae of HeLa cells expressing either Cidec-EGFP or Cidec untagged, identified by cryo-FM, were generated by cryo-FIB milling performed with a Scios DualBeam FIB/SEM (FEI) equipped with a Quorum stage (PP3010T) using a similar procedure as published before (21). Prior to milling, grids were coated with organometallic platinum using a gas injection system for 30s at 13mm working distance and 25° stage tilt. The electron beam was used for locating the cells of interest at 5 kV voltage and 13 pA current and for imaging to check progression of milling at 2 kV voltage and 13 pA current. Milling with the ion beam was performed stepwise. For rough milling the voltage was kept at 30 kV throughout all steps and milling was performed simultaneously from the top and the bottom of the lamella. The current and stage position were adjusted as follows: 1) 1 nA, 35° stage tilt until a lamella thickness of 20 *µ*m; 2) 0.5 nA, 25° tilt until 12 *µ*m; 3) 0.3 nA, 17° tilt until 3 *µ*m; and 4) 0.1 nA, 17° tilt until 1 *µ*m; 2). For the fine milling steps, the voltage was lowered to 16 kV and the current to 23 pA. The stage was first tilted to 18° and the lamella was milled only from the top, then the stage was tilted to 16° and the lamella was milled only from the bottom. These two steps resulted in a lamella thickness of approximately 0.5 *µ*m. Finally, the stage was tilted back to 17° and the lamella was milled simultaneously from the top and the bottom to a thickness below 0.3 *µ*m.

### Electron cryo-tomography (cryo-ET)

Cryo-ET data of FIB-milled cells was acquired on a Titan Krios microscope (Thermo Fisher Scientific) equipped with a K2 direct electron detector (Gatan) and a Quantum energy filter. SerialEM was used for acquisition (22). To identify the positions of the lamella, a montage of the central part of the grid was acquired at 171 nm pixel size in linear mode. Then a 5.1 nm pixel size montage of individual lamella were taken and used for finding interfaces between LDs. Tilt series were acquired from 0 ° to ± 60 ° with a dose-symmetric acquisition scheme (23), 1 ° increment in counting mode and a pixel size of 3.7 or 3.5 Å. Images were acquired in a tilt group size of 4, and 4 frames per tilt. Target defocus was set to −5 *µ*m. The dose per tilt series image was adjusted to 1 e^-^/A^2^ and the target dose rate at the detector was kept around 4 e^-^/px/s. Alignment of the tilt series frames was performed in IMOD using alignframes and the tilt series were aligned in IMOD using patch tracking (24, 25). Final tomograms were reconstructed at 7.4 or 7.1 Å pixel size using SIRT reconstruction with 10 iterations of a SIRT. For presentation in figure panels and movies, tomographic slices were filtered with a median filter.

### Analysis of distances between lipid droplets

Distance measurements were performed in cryoET tomograms by using a custom-made MATLAB (MathWorks) script. Control points along each monolayer were manually defined at positions where the monolayers are best visible. The membranes were traced every 7.5 or 7.1 Å in z direction through the tomographic volume. Based on these points, the surfaces of each monolayer are interpolated on a 2 nm regular grid. The distance between points of the interpolated surfaces are computed and various descriptive statistics computed. In particular, histogram of distances, minimum and maximum distance as well as the mean, median and the standard deviation of the distances were calculated. Each interface between a LD pair is considered as an individual event. Calculated medians of the distance at these interfaces were plotted as a scatter plot. The error bars represent mean +/-standard deviation.

### Determination of lipid droplet diameters

LD diameters shown in Figure 2g were measured in IMOD using the distance measurement tool. For the measurements, the 5.1 nm pixel size montage images of lamellae were used (see cryo-ET section). To confirm that the equatorial plane of individual LDs was roughly included, the corresponding reconstructed tomogram was inspected in a side view. LDs for which the equatorial plane was not part of the acquired volume were excluded from the analysis.

### Statistical tests

All P values reported in the text are based on statistical tests performed on the data presented in the corresponding Figure panels. In all applicable instances, we used the means of independent experiments to calculate P values, and plotted the corresponding data as ‘superplots’ (Fig. 1a, 1b, 3a, 3b and 3c) (26). All statistical tests and plots were done in Graphpad Prism. The following tests have been applied on the data shown in Fig. 1b: unpaired two-tailed t test; Fig. 2h: Ordinary one-way ANOVA, multiple comparisons; Fig. 2i: unpaired parametric two-tailed t-test; Fig. 3a: Welch ANOVA test (all constructs, +Dox), unpaired two-tailed t tests (EGFP +Dox vs. EGFP -Dox, and Cidec-SUMOstar +Dox vs. Cidec-SUMOstar -Dox) unpaired two-tailed t test with Welch’s correction (Cidec +Dox vs. Cidec -Dox); Fig. 3b: Welch ANOVA test (all constructs, +Dox), unpaired two-tailed t tests (EGFP +Dox vs. EGFP -Dox, Cidec-EGFP +Dox vs. Cidec-EGFP -Dox, Cidec-SUMOstar +Dox vs. Cidec-SUMOstar -Dox, and Cidec +Dox vs. Cidec -Dox); Fig. 3c: Welch ANOVA test; Fig. 5b: unpaired two-tailed t-test; Fig. 5e: nonparametric unpaired two-tailed Mann-Whitney t-test.

---

**Supplementary Figure S1:**
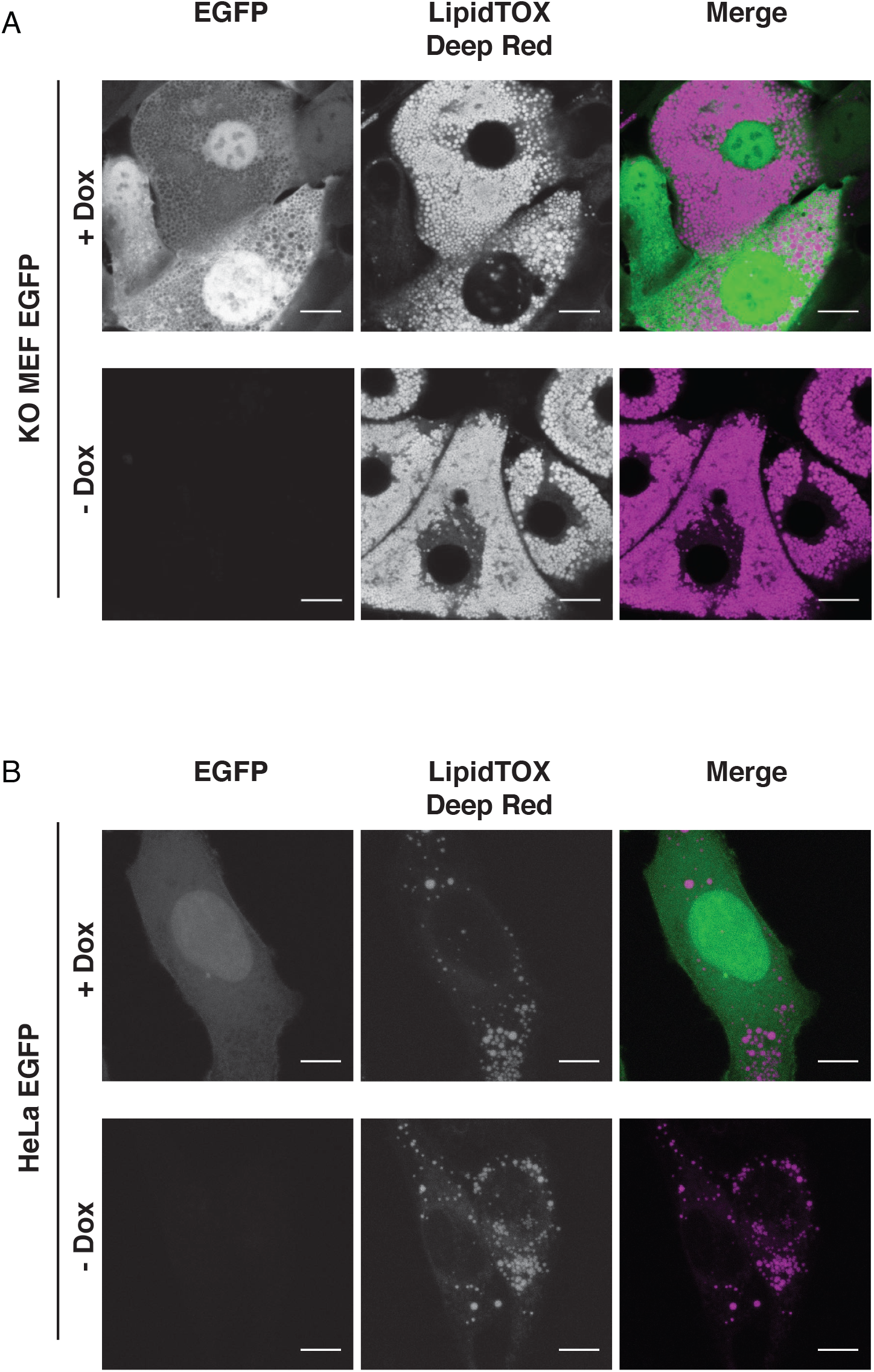
Expression of cytosolic EGFP. **A)** Fluorescence microscopy of fixed Cidec null MEF-derived adipocytes inducibly expressing cytosolic EGFP (green). LDs were stained with LipidTOX Deep Red dye (magenta). Cidec null MEF-derived adipocytes were induced with Doxycycline (upper panels, + Dox) or not induced (lower panels, -Dox) throughout the course of differentiation. Scale bars: 10 *µ*m. **B)** Fluorescence microscopy of a fixed HeLa cell line inducibly expressing cytosolic EGFP. LDs were stained with LipidTOX Deep Red. The upper panels represent HeLa cells induced with Doxycycline for cytosolic EGFP overexpression. The lower panels show the control cells in the absence of Doxycycline induction. All images included in the gallery are images of fixed cells taken 24 hours after the corresponding treatment (+ Dox/ -Dox). Scale bars: 10 *µ*m.

**Supplementary Figure S2:**
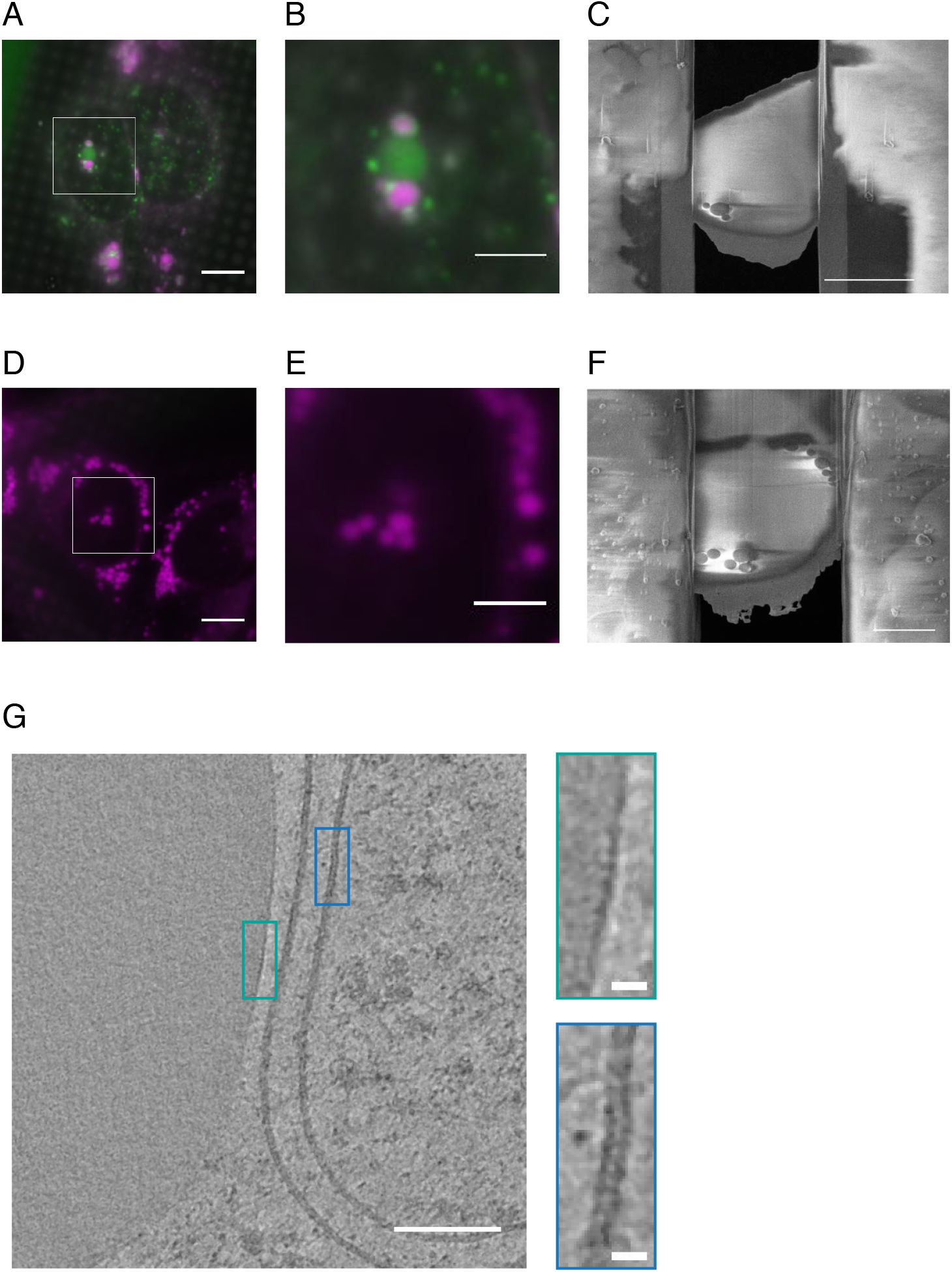
Cryo-CLEM of HeLa cells expressing Cidec constructs, representative examples. **A and B)** Cryo fluorescence microscopy of plunge-frozen HeLa cells inducibly expressing Cidec-EGFP, grown on a cryo-EM grid. Region for cryo-FIB milling was chosen based on LD size, indicating enlargement, and Cidec-EGFP enrichment at LD interfaces. **B)** Magnified view of the region shown in the white box in A. **C)** SEM overview image of a lamella generated from the HeLa cell shown in A and B, thinned by cryo-FIB milling. Interacting LDs identified by cryoFM are visible in the resulting lamella. **D and E)** Cryo fluorescence microscopy of plunge frozen-HeLa cells inducibly expressing untagged Cidec. Representative Hela cells grown on an EM grid. Region for cryo-FIB milling was chosen based on LD size and proximity. **E)** Magnified view of the region shown in a white box in A. **F)** SEM overview image of the lamella generated from the HeLa cell shown in D and E, thinned by cryo-FIB milling. Interacting LDs identified by cryoFM are visible in the resulting lamella. **G)** Comparison of the morphology of a monolayer with the morphology of a bilayer. Virtual slice through an electron-cryo tomogram of a HeLa cell inducibly expressing Cidec-EGFP. Green box shows a magnified view of a monolayer, blue box shows a magnified view of a bilayer. Scale bars: 10 *µ*m in A, C and D, 5 *µ*m in B, E and F, 100 nm in G large image, 10 nm in G magnified views.

**Supplementary Figure S3:**
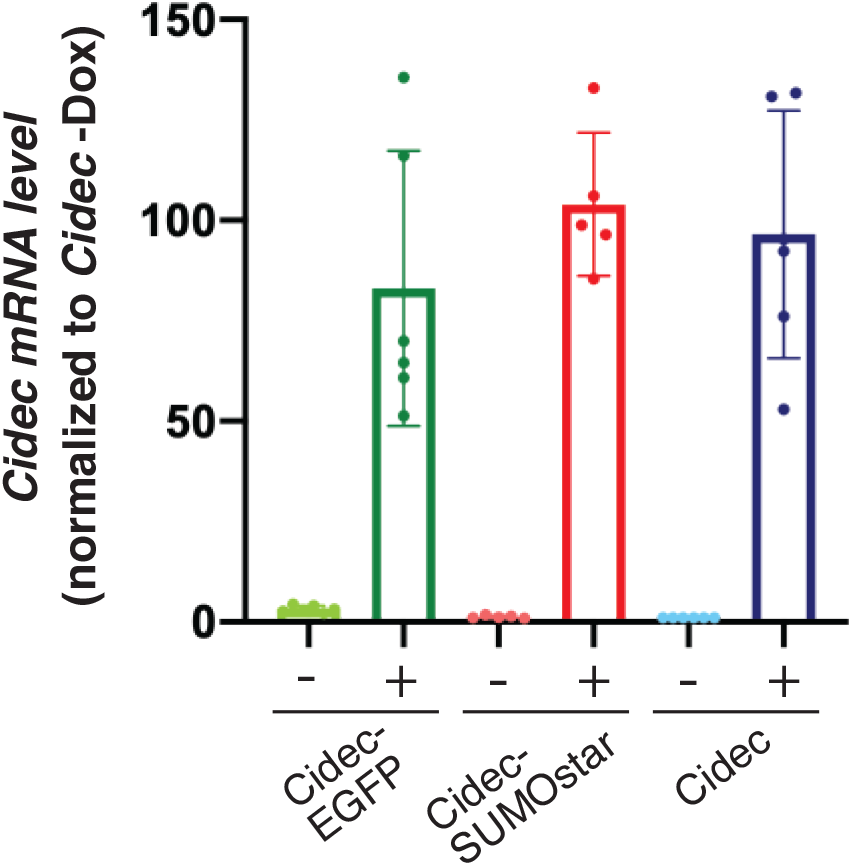
Expression levels of Cidec transcripts in HeLa cells transduced with the Doxycycline-inducible system. HeLa cell lines were induced with Doxycycline for Cidec expression in the presence of 200 *µ*M oleic acid for 24 hours. Comparable levels of Cidec transcripts were achieved by using 0.5 *µ*g/mL of Doxycycline in Cidec-EGFP and untagged Cidec HeLa cell lines and 2.0 *µ*g/mL of Doxycycline in the Cidec-SUMOstar HeLa cell line. mRNA levels were normalized to the level of untagged Cidec HeLa cell line without Doxycycline treatment. The plot shows mean Cidec mRNA expression levels with SD from at least 5 independent experiments.

## Supplementary movie legends

<u>Movies 1-4:</u>

Electron cryo-tomograms of HeLa cells expressing Cidec-EGFP, corresponding to Figures 2a (Movie 1), 2b (Movie 2), 2c (Movie 3), and 2d (Movie 4). The movie presents virtual slices along the z-axis of tomographic volume. The white bounding box indicates the dimensions of the tomographic volumes which are 1.43 *µ*m and 1.38 *µ*m in x and y, respectively.

<u>Movies 5-7:</u>

Electron cryo-tomograms of HeLa cells expressing untagged Cidec, corresponding to Figures 2e (Movie 5), 2f (Movie 6) and 2g (Movie 7). The movie presents virtual slices along the z-axis of tomographic volume. The white bounding boxes indicate the dimensions of the tomographic volumes which are 1.43 *µ*m and 1.38 *µ*m in x and y, respectively.

<u>Movies 8-10:</u>

Live FM of lipid droplets, stained with LipidTOX Deep Red, in cells expressing Cidec-EGFP (Movie 8), Cidec-SUMOstar (Movie 9) and untagged Cidec (Movie 10). Movie frames correspond to maximum projection images of z-stacks. Z-stacks were acquired at an interval of 2.5 min (Cidec-EGFP) or at intervals of 20 seconds (untagged Cidec and Cidec-SUMOstar). Imaging started 24h post-Doxycycline induction (Cidec-EGFP) or 2 h post-Doxycycline induction (untagged Cidec and Cidec-SUMOstar). Scale bars are 2 *µ*m.

